# Evidence that an HDAC2-targeted ASO persistently upregulates cortical acetylcholine and dopamine signaling via a CREB | Gs positive feedback loop

**DOI:** 10.1101/2020.09.24.312264

**Authors:** Teresa H. Sanders

## Abstract

Epigenetic modulation of neural circuits facilitates learning and memory. Here, we examined specific inhibition of histone deacetylase 2 (*HDAC2*) expression in rats receiving a single intracerebroventricular injection of *HDAC2*-targeted anti-sense oligonucleotides (ASOs) one month prior to cognitive testing. The *HDAC2* ASO-injected rats displayed increased novelty preference, decreased cortical and hippocampal *HDAC2* mRNA and protein, and upregulated gene expression that persisted 1-month post-injection. Cortical RNA-seq revealed strongly increased transcription of a subset of cyclic adenosine monophosphate (cAMP)-response element binding (*CREB*) genes known to influence synaptic plasticity, along with dopamine (*DRD1, DRD2*) and adenosine (*ADORA2A*) G-protein-coupled receptors (GPCRs). Our analysis identified evidence of a positive-feedback loop that amplified expression of CREB-regulated Gs GPCRs and genes in cAMP/Gs/Gi signaling pathways. Additionally, we found differential expression of enzymes that shift neurotransmitter biosynthesis away from norepinephrine and toward dopamine and acetylcholine (DBH, CHAT). We also observed increased expression of genes important for neurotransmitter packaging (*SV2C, VMAT*) and release (*SYT9*). The data indicate that persistent inhibition of *HDAC2* expression enables long-lived enhancement of aspects of cognition through increased cortical transcription of a subset of CREB-regulated genes amplified by a positive-feedback mechanism that increases synaptic plasticity and shifts neurotransmitter balance toward increased dopaminergic and cholinergic signaling.

## Introduction

Long-term memory requires transcriptional changes that are facilitated by epigenetic modulation of DNA accessibility (Korzus et al., 2004; Miller et al., 2008). To promote the necessary transcriptional changes, DNA that is tightly wrapped around histones and wound into heterochromatin must be made more accessible. Acetylation of lysines within the N-terminal tail of histones relaxes chromatin compaction and facilitates transcription (Zentner et al., 2013). Accordingly, studies have shown that histone acetylation and deacetylation are fundamental epigenetic modulations linked to cognition (Alarcon et al., 2004; Korzus et al., 2004; Miller et al., 2008; Sanders et al., 2019). Enzymes known as histone acetyltransferases (HATs), such as those found in CREB-binding proteins (CBPs), can activate transcription by transferring acetyl groups to histones. On the other hand, histone deacetylase enzymes, or HDACs, facilitate chromatin compaction by removing acetyl groups, thereby repressing transcription.

Decreasing acetylation by inhibiting HATs impairs long-term memory (Alarcon et al. 2004; Wood et al. 2005), while increasing acetylation by inhibiting HDACs has been shown to enhance hippocampal long-term potentiation (Vecsey et al., 2007), memory (Levenson et al. 2004; Hawk et al. 2011; Itzhak et al., 2013), neuronal development (Cho and Cavalli, 2014), and cognition (Penney et al., 2014). In mice, loss of one of the eleven HDAC isoforms, HDAC2, has also been shown to improve performance on hippocampal and prefrontal-cortex dependent learning tasks (Guan et al. 2009; Morris et al. 2013). *HDAC2* has been found to be overexpressed in Alzheimer’s disease (AD) in humans and rodents, and its inhibition has been shown to reduce signs of AD in a mouse model (Graff et al. 2012). Accordingly, specific inhibition of *HDAC2* has been a goal of pharmacological design (Wang et al. 2005; Choubey and Jeyakanthan 2018).

Antisense oligonucleotides (ASOs) are clinically useful for treating disease (Alter et al., 2006; Downes et al., 2006; DeVos et al., 2013; Stein and Castanotto 2017) since they provide specificity through base pairing with a target messenger RNA (mRNA). The ASOs used in this study target *HDAC2* mRNA. Previously, these ASOs have been shown to enhance memory in wild-type mice in object location memory tests and to rescue impaired memory in a mouse model of autism (Kennedy et al. 2016). In mice, a single *HDAC2* ASO injection reduced *HDAC2* expression for over 4 months, and increased object location memory for 8 weeks (Poplawski et al., 2020). ASOs targeting other mRNAs have been shown to reduce expression of their target genes in the central nervous system for months after delivery of the drug (Kordasiewicz et al. 2012; Southwell et al., 2014; Meng et al. 2015). ASOs can act through recruitment of RNaseH1 to the RNA/ASO hybrid and subsequent degradation of the RNA (Wu et al. 2004) or by modulation of splicing (Merkhofer et al. 2014). Recent studies have also shown evidence that ASOs may interfere directly with transcription of the target gene (Poplawski et al., 2020).

Although HDAC inhibitors have been shown to enhance memory processes by activation of genes regulated by the CREB: CBP transcriptional complex (Vecsey et al., 2007; Guan et al., 2009), the specific mechanisms linking long-term ASO inhibition of *HDAC2*, cortical plasticity, and cognitive enhancements are poorly understood. Putative mechanisms include those that modulate norepinephrine, dopamine, and acetylcholine since these neurotransmitters are known to play important roles in learning and memory (Rasmusson et al., 2000; Myhrer et al., 2003; Berridge and Waterhouse, 2003; Sara, 2009) and are pharmacologic targets for neurodegenerative disorders such as AD (Hardy et al., 1985; Chalermpalanupap et al., 2013) and Parkinson’s disease (Cools et al., 2001; Mattay et al., 2002; Verschuur et al., 2019). Evidence for the importance of their role in the brain is emphasized by recent projection-tracing studies that have revealed the widespread nature of neurotransmitter-releasing neuronal circuits (Bjorklund and Dunnett, 2007; Chandler et al., 2014; Aston-Jones and Waterhouse, 2016). Here, we investigate gene expression alterations following a multi-day novelty preference task to discover that acetylcholine and catecholamine neurotransmitter pathway modulation enabled by a CREB | Gs signaling positive feedback loop, along with upregulated expression of synaptic plasticity, cellular differentiation, and forebrain development genes, underlies persistent improved cognition in rats injected with *HDAC2* ASOs.

## Results

### ASO-injected rats display long-lived reductions in *HDAC2* mRNA and protein

One month after intracerebroventricular (ICV) injection of *HDAC2* ASOs, protein (Western Blot) and transcripts (qPCR) from cortical and hippocampal tissues were quantified (Fig. 1). Since these initial results showed *HDAC2* mRNA and protein were reduced more after ASO1 injections compared to ASO2 (Fig. 1C), experimentation and analysis proceeded with ASO1. Rats receiving *HDAC2* ASOs had significantly reduced *HDAC2* expression compared to rats receiving ICV injections of saline or non-targeting ASO (Fig. 1C, ASO1 cortex: log2(fold-change) = −1.62, −1.40, p < 0.01, mRNA and protein respectively, ASO1 hippocampus: log2(fold-change) = −2.33, −1.85, p < 0.01, mRNA and protein respectively).

**Figure 1.**
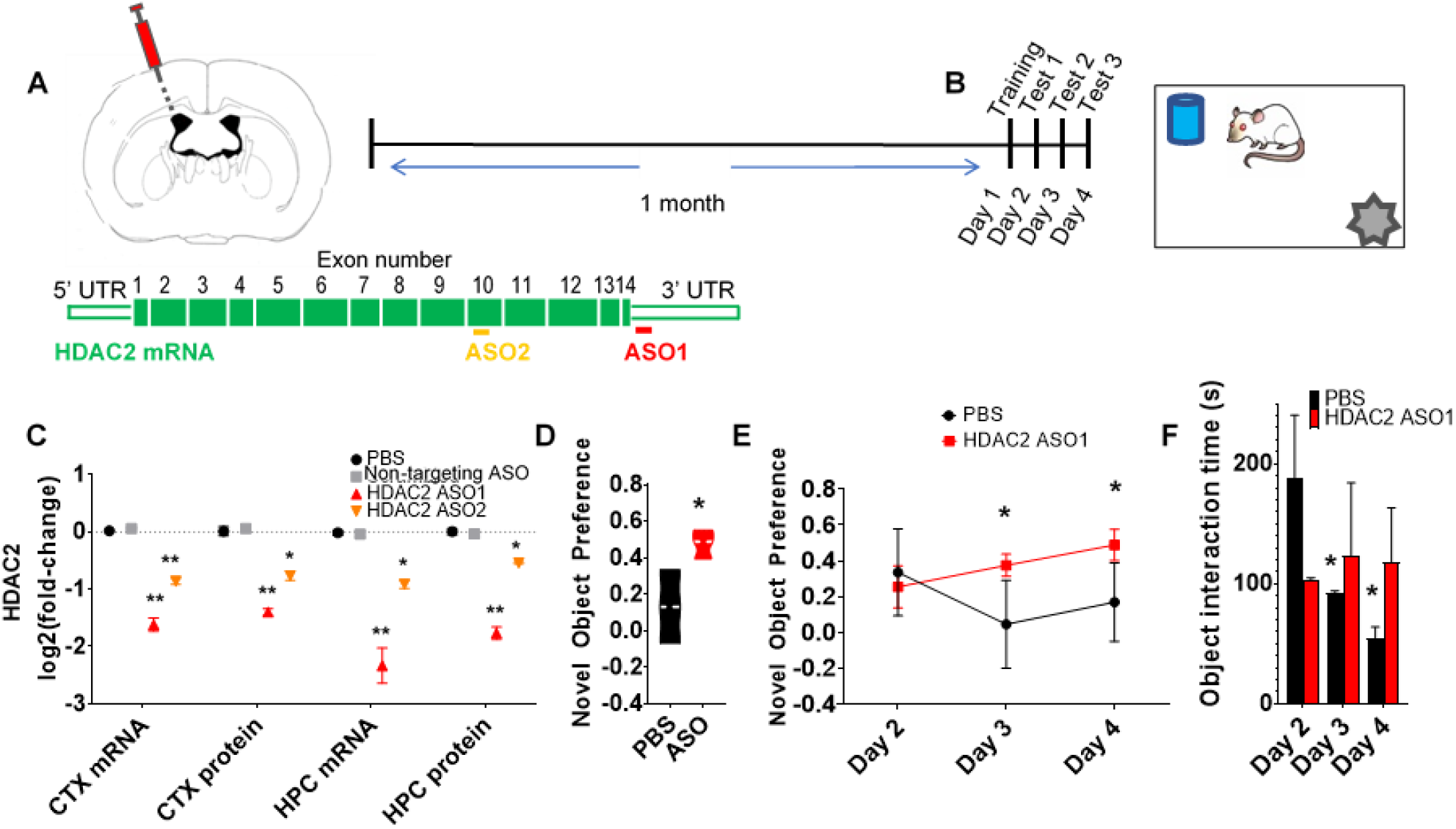
A single injection of *HDAC2* ASO results in enhanced novelty preference and long-lived reduction of *HDAC2* expression. (**A**) *HDAC2* ASOs were delivered to rats via intracerebroventricular (ICV) injection, (**B**) A multi-day novel object preference test was administered one month post-injection, (**C**) Cortical (CTX) and hippocampal (HPC) tissues were extracted immediately following the last session of behavioral testing to confirm reduced *HDAC2* protein and mRNA in hippocampus and cortex of rats injected with one of two *HDAC2* ASOs (ASO1, ASO2) compared to rats injected with the same volume of PBS or a non-targeting ASO (NTA), (**D**) Overall novel object preference was increased in ASO1-injected rats compared to rats injected with the same volume of PBS. (**E**) Novel object preference was similar in PBS and *HDAC2* ASO1 groups for the first day of testing, but ASO1-injected rats showed greater preference for the novel objects on the last two days of testing, (**F**) PBS-injected controls interacted with the objects less on each consecutive day, while ASO1-injected rats showed the same or greater object interaction on consecutive days. Behavioral tests were performed on rats receiving either PBS or *HDAC2* ASO1 (N = 3 rats per category, later used for RNA-seq analysis), qPCR tests were performed on a separate cohort of rats with N = 3 rats per category (PBS, NTA, *HDAC2* ASO1, *HDAC2* ASO2, N = 12 rats total). SEM error bars, * p ≤ 0.05, ** p ≤ 0.01.

### *HDAC2* ASO1 injection improves performance on a novel object recognition task

Novel object recognition tests are commonly used to assess cognition in rodents (Broadbent et al., 2010; Antunes and Biala, 2012). Previously, we reported that electrical stimulation administered through implanted cuff electrodes on the left cervical vagus nerve (VNS) in Sprague-Dawley rats during 30 minutes of object familiarization improved next-day performance on a novel object recognition task (Sanders et al., 2019). Performance was assessed by calculating the difference in the amount of time spent interacting with a novel object (t_o2_) compared to the familiar object (t_o1_), expressed as a fraction of the total time spent interacting with either object 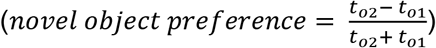. The test was administered for 10 minutes on 3 consecutive days with new objects rotated in each day (see methods, Fig. 1B). In the current study, *HDAC2* ASO1 injected Sprague-Dawley rats tested with this paradigm 1-month also preferred the novel object more strongly than did saline-injected controls (Δ_novel_object_preference_ = 0.35, p = 0.03, n = 6, Fig. 1D).

Although the overall cumulative novel object preference was significantly increased for *HDAC2* ASO-injected rats, there was no difference in novelty preference between sham and treated rats on the initial day of testing (Fig. 1E). Investigation of the time spent interacting with objects each day further revealed that, for PBS-injected rats, object interaction time was diminished on consecutive days of exploration (reduced 51%, 41%, Days 3 and 4 respectively, compared to the previous day, p = 0.05). This trend toward decreasing novelty exploration over time has been observed in previous studies (Gaskin et al., 2010; Sanders et al., 2019). However, in the present study, *HDAC2* ASO1-injected rats interacted with the objects for a similar amount of time each day (Fig. 1F), with novelty preference peaking on the last test day (Fig. 1E, *novel object preference* = 0.48, p = 0.03).

### *HDAC2* ASO1 injection modulates cortical plasticity gene expression

Analysis of RNA-seq results from hippocampal-adjacent cortex from ASO1-injected rats (Fig. 2) revealed evidence of gene expression representative of both glial (*OLIG1, OLIG2, GFAP*) and neuronal (*MAP2, SYP*) cells. However, no significant differences were found between the PBS-injected control and ASO1-injected rat cell identity transcripts (Fig. 2B). Consistent with the qPCR results, RNA-seq results showed a significant decrease in cortical *HDAC2* expression in ASO1-injected rats (Fig. 2D, log2(fold-change) = −0.9, p = 2.0e-3).

**Figure 2.**
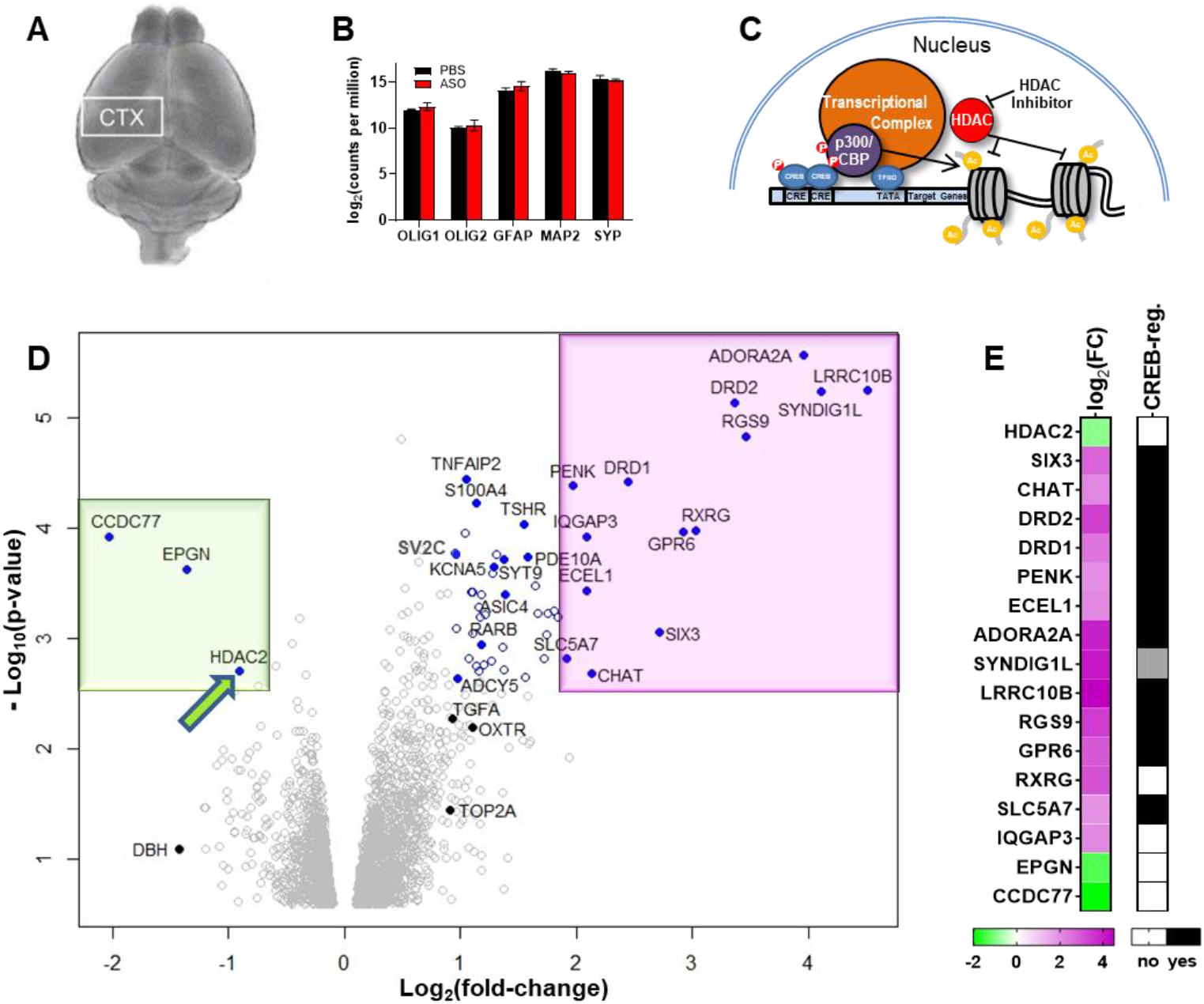
HDAC2 ASO injection increases cortical transcription. (**A**) RNA was extracted from hippocampal-adjacent cortical tissues. (**B**) Cell identity transcripts were not significantly changed between ASO and PBS cohorts. (**C**) HDAC inhibitors have previously been found to increase transcription of genes with cAMP-responsive element (CRE) binding domains. (**D**) RNA-seq confirmed reduced HDAC2 transcripts and increased transcription of many genes including a set of highly upregulated genes (purple region). Blue circles indicate differentially expressed genes with FDR < 0.3. (**E**) Genes with the most significantly changed transcription, ordered by magnitude of correlation with HDAC2, were primarily genes known to be CREB-regulated. N = 3 rats per category (same cohort used for behavioral testing)

Overall, cortical transcripts were increased for 516 genes and decreased for 184 genes (p < 0.05). The counts per million (cpm) for genes with the largest increases correlated inversely with the *HDAC2* cpm (Fig. 2D-E (purple), magnitude of the correlation coefficient, ∥*ρ*∥ ≥ 0.74, p < 0.01) and included genes that encode LRRC10B (log_2_(FC) = 4.5, p = 5.7e-6, leucine rich repeat containing 10B, localized to the nucleoplasm), SYNDIG1L (4.1, p = 5.8e-6), ADORA2A (4.0, p = 2.7e-6, Adenosine A2A (Gs) receptor, regulates glutamate and dopamine release) (Hack et al., 2003, Morelli et al., 2007), RGS9 (3.5, p = 1.5e-5), DRD2 (3.4, p = 7.3e-6, D2 dopamine receptor (Gi-protein coupled), found to be important for cognitive flexibility) (Cameron et al., 2018), RXRG (3.0, p = 0.002), GPR6 (2.9, p = 0.0001), SIX3 (2.7, p = 0.0009), DRD1 (2.4, p = 3.8e-5, D1 dopamine receptor (Gs), regulates neuronal growth and development, mediates behavioral responses and memory, found to be important for cognitive stability) (Cameron et al., 2018), IQGAP3 (2.1, p = 0.0001), ECEL1 (2.1, p = 0.0004), CHAT (choline acetyltransferase), PENK (2.0, p = 4.1e-5, proenkephalin), and SLC5A7 (1.9, p = 0.001, choline transporter).

Analysis of this top upregulated group of differentially expressed (DE) genes (log_2_(FC) > 1.9, p < 0.0021) revealed that the majority either contain a CRE domain or respond to enhancers with CRE domains (Figs. 2C, 2E). CRE-binding protein (CREB) transcriptional regulation is important for neuronal plasticity and long-term memory formation in the brain and has been shown to be integral in the formation of spatial memory (Silva et al., 1998) while CREB downregulation has been observed in the pathology of AD (Pugazhenthi et al., 2011).

Although increased CRE regulation was clearly associated with the top group of upregulated genes (Fig. 2D-E), not all these highly upregulated genes contained known CRE domains. Top DE genes without confirmed CREB-regulation in rats include: *RXRG*, that forms heterodimers with retinoic acid, thyroid hormone, and vitamin D receptors to facilitate both DNA binding and transcriptional function on their respective response elements, *IQGAP3*, that expresses a calmodulin-binding scaffolding protein that plays a role in neuronal morphogenesis such as neurite outgrowth (Wang et al., 2007), and *SYNDIG1L*, Synapse Differentiation Inducing 1 like gene (Fig. 2E, gray, although known to be affected by an enhancer regulated by CREB in humans (Zhang et al., 2005), not yet confirmed to be CREB-regulated in rats (Impey et al., 2004)).

The DE genes in this study with potential CREB-regulation were not limited to the top DE genes. Many with smaller fold-changes also contain promoter CRE domains or interact with enhancers containing CRE domains. Additionally, not all genes with a CRE domain were upregulated (e.g. *BDNF* and tyrosine hydroxylase (*TH*) were not upregulated, although both have previously been found to contain one or more CRE motifs and to respond to associated HAT activity).

### *HDAC2* inhibition with *HDAC2* ASOs alters expression of key enzymes in catecholamine and acetylcholine biosynthesis pathways

The RNA-seq results implicate the effects of increased CREB binding protein HAT activity in the ASO-injected rats (Fig. 2D-E). However, although increased transcription of *DRD1* and *DRD2* suggests modulation of dopamine (DA), the overall effect on dopamine signaling cannot be inferred from the receptor transcriptional changes. To gain further insight into the biochemical pathway changes induced by *HDAC2* inhibition (*HDAC2*i) → CREB → DA, expression of the enzymes involved in the dopamine biosynthesis pathway was examined (Fig. 3A). Since acetylcholine is another important neurotransmitter involved in cognition, the enzymes that control acetylcholine availability were also examined (Fig. 3B). Expression of the primary acetyl choline biosynthesis enzyme, choline acetyltransferase (*CHAT*), correlated strongly with the decrease in *HDAC2* expression and appeared in the group of most strongly upregulated genes (Fig. 2E). Further examination revealed that, among neurotransmitter biosynthesis enzymes, only *CHAT* and dopamine β hydroxylase (*DBH*) were differentially expressed (log_2_(FC) = 2.1 and log_2_(FC) = −1.6, respectively, Fig. 3C). These two enzymes were also the only biosynthesis pathway enzymes found to be related to the *HDAC2* inhibition. *DBH* and *HDAC2* expression correlated strongly (ρ = 0.89, p = 0.02, Fig. 3C-D). Reduced DBH results in less conversion of DA to norepinephrine (NE) during catecholamine biosynthesis (Fig. 3A). The lack of change in other enzymes in the pathway indicates that it is likely that more DA (and less NE) was available in the cortex of *HDAC2*-inhibited rats (Devoto et al., 2015). The increased expression of dopamine receptors DRD1 and DRD2 further suggests that the increased DA may have had increased interaction with postsynaptic neurons, thus potentially increasing the effects of DA on cognition (Cameron et al., 2018).

**Figure 3.**
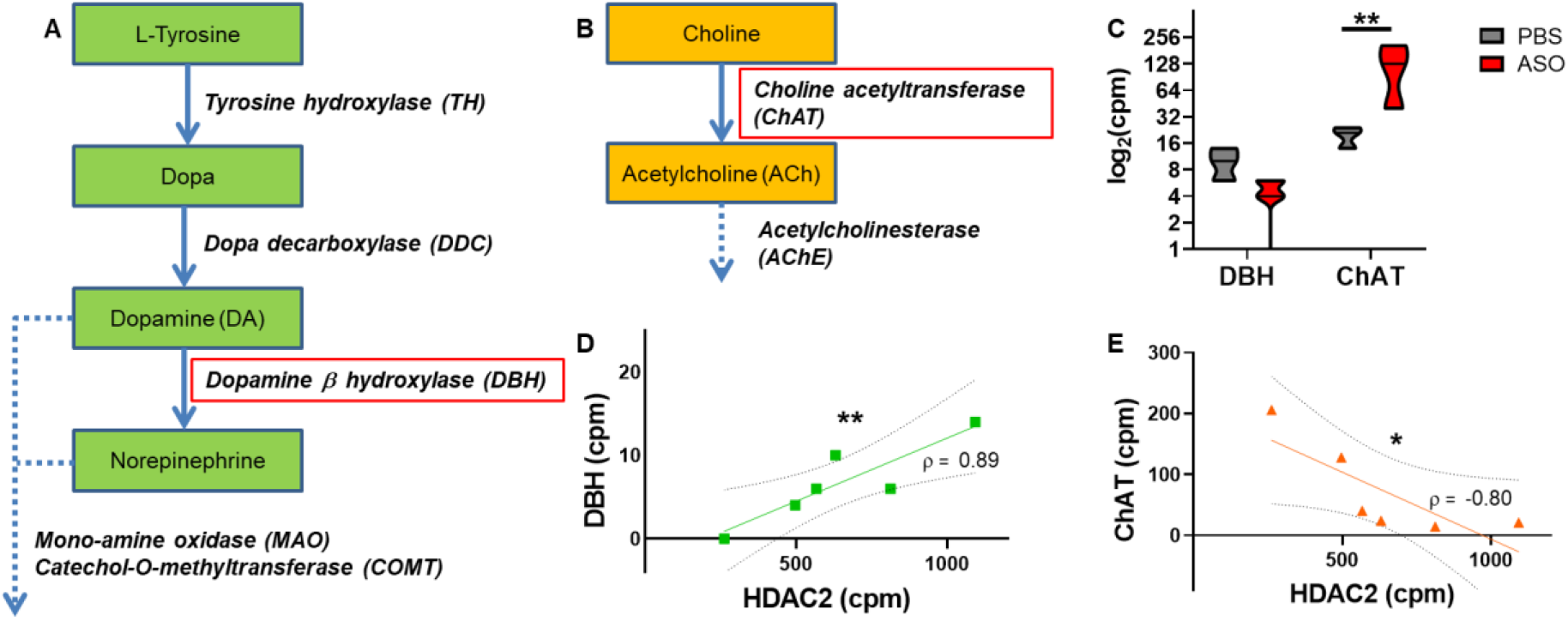
Key neurotransmitter biosynthesis enzymes display altered expression after long-term *HDAC2* inhibition. (**A**) Expression of dopamine beta hydroxylase (*DBH*), the gene that encodes the enzyme responsible for converting dopamine to norepinephrine was reduced in *HDAC2* ASO-injected rats, while (**B**) expression of choline acetyltransferase (*CHAT*), the gene that encodes the enzyme responsible for converting choline to acetylcholine, was increased. (**C**) The fold-change in *CHAT* expression was larger than the change in *DBH* expression. (**D**) However, the magnitude of the correlation between *DBH* and *HDAC2* counts per million (cpm) (ρ = 0.89) was greater than (**E**) the magnitude of the inverse correlation between *CHAT* and *HDAC2* cpm (ρ = −0.80). Curved lines show 95% confidence intervals. * p < 0.05, **p < 0.01.

Similarly, while *CHAT* expression was increased and correlated significantly with reduced *HDAC2* (ρ = −0.8, p = 0.05, Fig. 3E), Expression of acetylcholinesterase (*AChE*), the gene that encodes the enzyme that catalyzes the breakdown of acetylcholine, was not significantly changed or correlated, suggesting a pathway shift resulting in increased available acetylcholine in *HDAC2*-inhibited rats (Fig. 3B). Taken together, these results indicate that *HDAC2* inhibition modulates-pathway enzyme transcription toward increased dopamine and acetylcholine, and decreased norepinephrine.

### *HDAC2*i increases transcripts associated with CREB-activation, neuronal synaptic activity and re-organization, cellular signaling and differentiation, and forebrain development

Gene ontology analysis of the RNA-seq results identified significantly increased biological processes (Fig. 4), summarized here in 4 categories: CREB-activated transcription (Fig 4A), neuronal synaptic activity and organization (Fig. 4A-C), cellular signaling and differentiation (Fig. 4B), and forebrain development (Fig. 4B). Fold-changes (FC) that have not been previously reported will be stated here as (*log_2_(FC)*, p = *p-value*).

**Figure 4.**
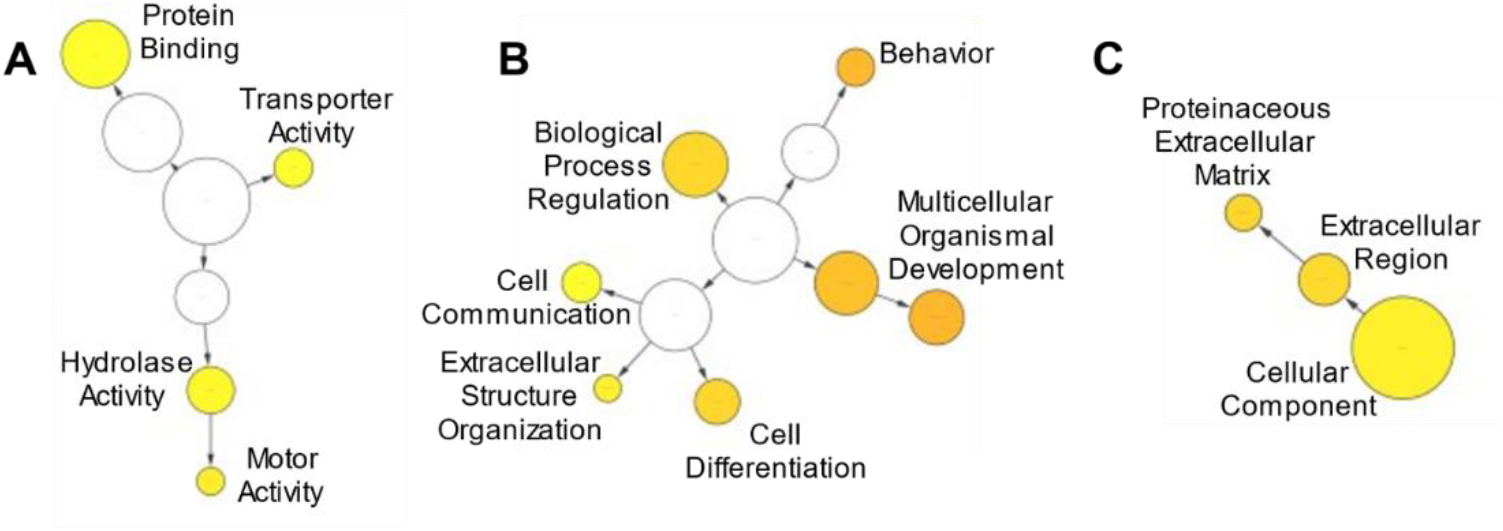
Gene ontology (GO) analysis reveals upregulation of multiple biological process gene sets. (**A**) Upregulated gene sets included HDAC/CREB-associated DNA-protein binding and hydrolase effector genes, along with CREB-associated genes known to facilitate motor activity, (**B**) cellular differentiation and communication genes related to developmental and behavioral changes, and (**C**) genes encoding synaptic and membrane-associated proteins that drive extracellular reorganization. Circle size indicates number of genes changed within the GO category. Circle color indicates the significance of the changes (**white → orange** corresponds to **least → most** significant).

#### CREB-activated transcription

The cAMP response element binding protein (CREB) is a nuclear transcription factor regulated by phosphorylation (Fig. 2C) via protein kinase A (PKA). PKA is, in turn, activated by cAMP (cAMP→PKA→CREB_phosphorylation). Many of the upregulated DE genes, including most of the top DE genes, were found to be CREB-regulated. Several of these genes were related to G-protein coupled receptor (GPCR) signaling, many of which, in turn, regulate cAMP by modulating its catalyzing enzyme, adenylyl cyclase (AC). RNA-seq results confirmed that AC expression was upregulated (*ADCY5*, 1.0, p = 0.002), revealing further evidence that a positive feedback loop between CREB, GPCRs, and cAMP signaling led to the strongly increased set of CREB-regulated Gs GPCR signaling transcripts. Examination of the top DE genes (False Discovery Rate, FDR < 0.3) revealed that none are Gq GPCRs. This is somewhat surprising since 1) the observed transcriptional changes suggest increased cholinergic activity and 2) CNS muscarinic acetylcholine receptors act primarily through Gq signaling (*CHRM1*). However, note that Gq associated intracellular signaling does not facilitate a positive feedback loop involving cAMP/PKA since Gq GPCR effects are mediated by phospholipase C (Fig. 5A). On the other hand, all four of the GPCRs in the top DE genes are adenylyl cyclase linked (Gs/Gi). This evidence combined with the observed increases in AC and CREB binding protein expression implicate Gs GPCRs as players in the putative GPCR → adenylyl cyclase → cAMP → PKA → CREB positive feedback loop that amplified expression of Gs GPCRs (Fig. 5B).

**Figure 5.**
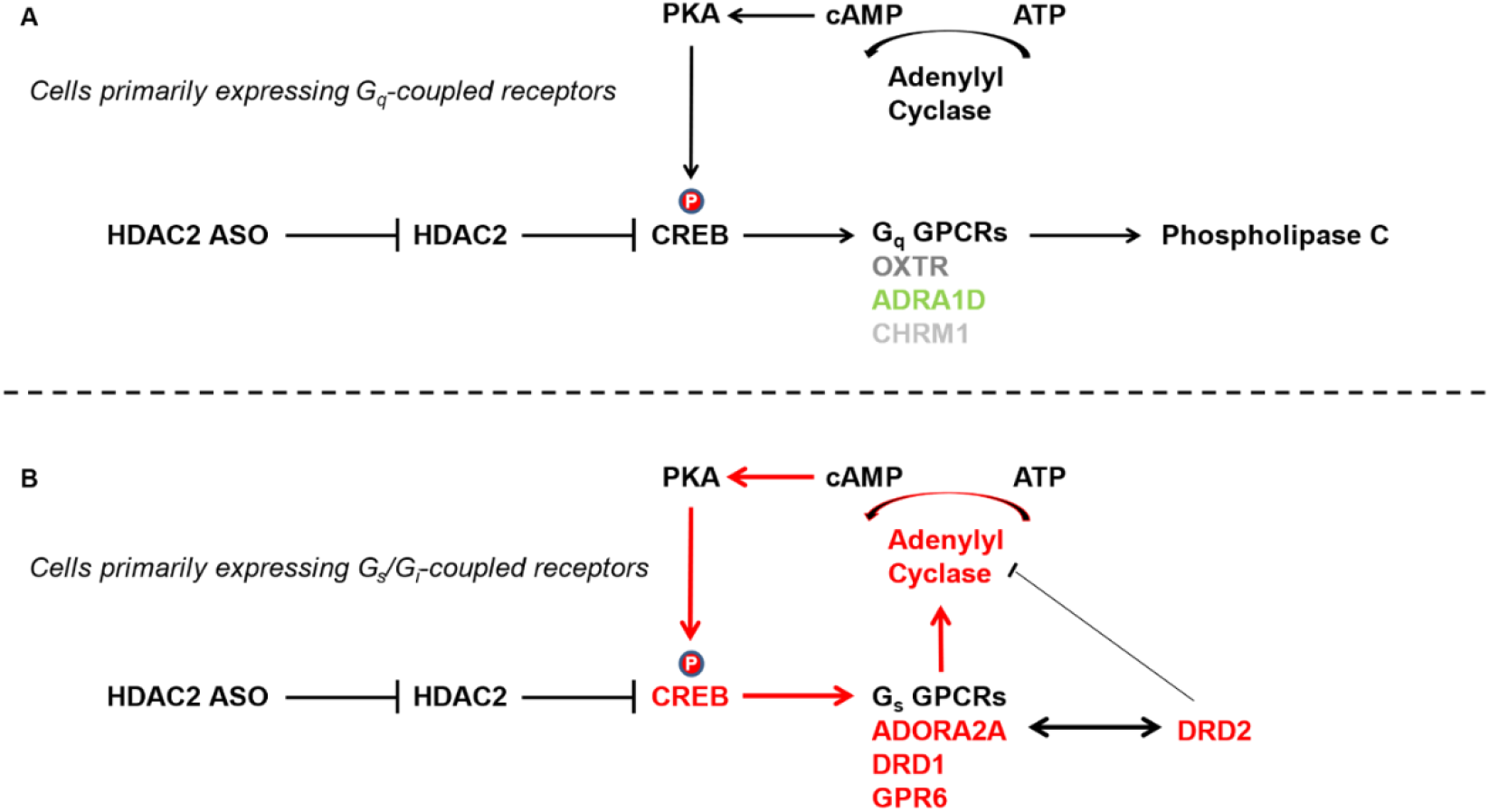
Gs, but not Gq, GPCRs enable an *HDAC2*i-induced CREB / G-protein-signaling positive feedback loop. (**A**) Gq receptors, expressed by genes such as *OXTR* (oxytocin receptor, log_2_(FC) = 1.1, p = 0.006)), *ADRA1D* (adrenergic receptor, log_2_(FC) = −0.45, p = 0.05), and *CHRM1* (muscarinic acetylcholine receptor, unchanged) primarily act by upregulating phospholipase C. (**B**) Gs receptors, expressed by genes such as *ADORA2A* (adenosine receptor, log_2_(FC) = 4.0, p = 2.7e-6), *DRD1* (dopamine receptor, log_2_(FC) = 2.4, p = 3.8e-5), and *GPR6* (log_2_(FC) = 2.9, p = 0.0001) upregulate cyclic AMP (cAMP) by increasing adenylyl cyclase. cAMP, in turn, activates protein kinase A (PKA) which phosphorylates CREB, thus creating a positive feedback loop that further promotes CREB-regulated expression. Note that the expression of CREB-regulated Gs receptor genes were orders of magnitudes larger than those for CREB-regulated Gq receptor gene *OXTR*.

#### Neuronal Synaptic Activity and Organization

Top DE G-protein signaling genes related to synaptic activity and organization included Gs GPCRs *ADORA2A, DRD1, DRD2*, and *GPR6* (upregulates cAMP and promotes neurite outgrowth), along with *RGS9*, which expresses a protein that modulates G proteins by promoting their deactivation and regulates dopamine and opioid signaling in the brain (Rahman et al., 2003). Mice deficient in RGS9 exhibit motor and cognitive difficulties (Blundell et al., 2008). These genes are all known to be CREB-regulated (Fig. 2D-E). Other G-protein signaling-related DE genes included the gene that encodes Gq GPCR oxytocin receptor (*OXTR*, log_2_(FC) = 1.1, p = 0.006, also CREB-regulated). Further top DE genes encode proteins important for synaptic activity and organization: *CHAT* (acetylcholine synthesis enzyme) and *SLC5A7* (aka *CHT*, choline transporter, CREB-regulated), *SYNDIG1L* (excitatory synapse regulator, Kalashnikova et al., 2010), *SLC35D3* (facilitates D1R emergence from endoplasmic reticulum, 1.7, p = 0.0006), and *IQGAP3* (neuronal morphogenesis regulator). Additional DE genes encoding proteins important for synaptic activity and organization included genes that encode cation channel with high affinity for sodium, *ASIC4* (1.4, p = 0.0004), calcium sensor / regulator of neurotransmitter release, synaptotagmin, *SYT9* (1.38, p = 0.0019), synapse excitability regulating voltage-gated potassium channel, *KCNA5* (1.29, p = 0.0002, CREB-regulated), regulator of neurite outgrowth, *SLITRK6* (1.0, p = 0.006), and vesicular proteins, *SV2C* (0.96, p = 0.00017). These neuronal synaptic activity and organization expression changes, together with the shifts in neurotransmitter biosynthesis enzyme pathways, support increased dopaminergic (Fig. 6A-B) and cholinergic neuronal signaling (Fig. 6C-D). Note that, in the cortex, some terminals co-release dopamine and norepinephrine, and re-uptake of both occurs primarily through the norepinephrine transporter (NET). However, investigation revealed that neither dopamine transporter (*DAT*) or *NET* transcripts were significantly changed, suggesting increased dopamine in the synaptic cleft (Fig. 6B). Finally, although the identity of the affected post-synaptic neurons cannot be definitively determined from the data (nor the identity of potential additional pre-synaptic neurons, or neurons impacted by volume transmission), the large increase in expression of *SYNDIG1L*, known to regulate excitatory synapses, suggests post-synaptic and/or downstream involvement of glutamatergic neurons.

**Figure 6.**
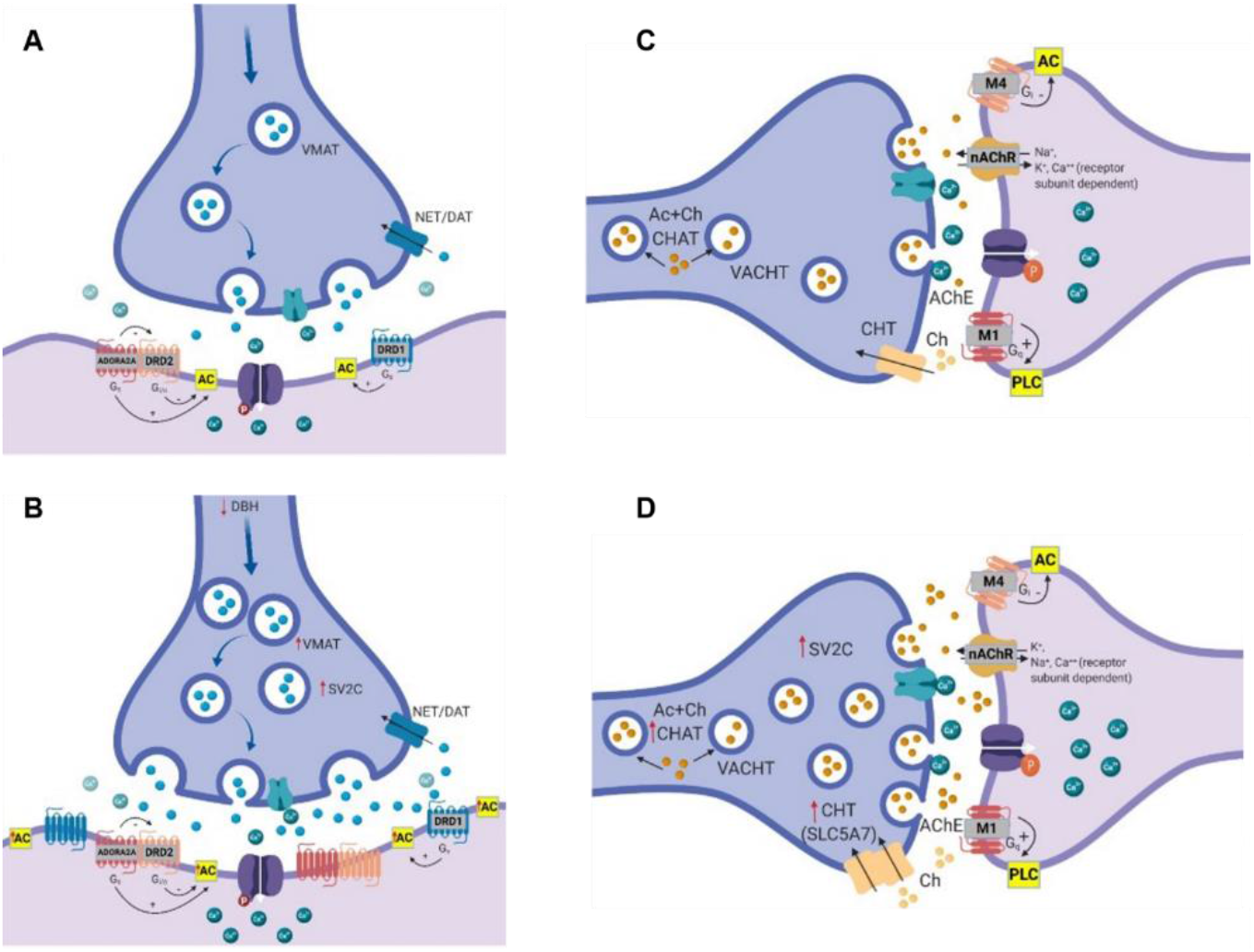
Synaptic signaling gene expression is dramatically upregulated 1 month after *HDAC2* inhibition. (**A**) Simplified diagram depicting putative pre- and post-synaptic neurons in control and (**B**) *HDAC2*-inhibited (*HDAC2*i) rats. *HDAC2*i induced changes favorable to dopamine (DA) biosynthesis (*DBH*) and increased vesicular packaging (*VMAT* and *SV2C*), as well as G protein-coupled receptor (GPCR) expression (*DRD1, DRD2, ADORA2A*). Increased cAMP signaling was implicated by upregulation of its synthesizing enzyme, adenyl cyclase (*AC/ADCY*). (**C**) Cholinergic signaling indicated by differential gene expression in control and (**D**) *HDAC2*i rats pointed to increased acetylcholine production (*CHAT*), vesicular packaging (*SV2C*), and choline transporter uptake (*CHT/SLC5A7*). Increased synaptotagmin (*SYT9*, not pictured) further suggests increased neurotransmitter release in (**B&D**). Upregulated AC and GPCRs are indicated by increased icon numbers. Other genes with upregulated expression are indicated by red arrows in (**B**) and (**D**). FDR < 0.3

### *HDAC2* expression and broad transcriptome changes correlate with the total time spent interacting with objects

Surprisingly, despite increased novel object preference in the *HDAC2* ASO-injected rats, we did not find correlations between novel object preference and expression of individual genes. Instead, strong correlation was observed between the time spent exploring the objects and the transcriptome changes (Supp. Fig. 1A). 882 genes, including *HDAC2*, correlated with object interaction time after the initial day of testing (Behavior Days 3-4, ∥*ρ*∥ ≥ 0.7, p < 0.05).

Expression of neurotransmitter biosynthesis pathway enzyme genes *CHAT* and *DBH* (Fig. 3) correlated significantly with object interaction time (Supp. Fig. 1B-C). Intra-correlation analysis of the most increased DE genes (FDR < 0.3) that correlated with object exploration time (∥*ρ*∥ ≥ 0.7, p < 0.05), and genes for associated synaptic proteins and enzymes that correlated strongly with object exploration time (∥*ρ*∥ ≥ 0.8, p < 0.01) revealed 4 gene clusters (Supp. Fig. 1D). The first cluster contained mainly CREB-regulated DE genes outside the top group of DE genes. The second group contained CREB binding protein (*CREBBP*) and *HDAC8*. The third cluster contained the top group of CREB-responsive GPCR genes (*ADORA2A, DRD1, DRD2*, and *GPR6*) along with adenylyl cyclase (*ADCY5*) and *CHAT*. The final group contained DE genes associated with cell differentiation (*RXRG, SIX3, CRABP1*) and synaptic organization. Other findings of interest include the high similarity in gene cross-correlation patterns for frequent heterodimer partners DRD2 and ADORA2A (correlation between expression of *DRD2* and *ADORA2A* = 0.999, p = 1.5e-12), the correlation of expression of pre-synaptic *α* adrenergic Gi GPCRs with object interaction time (*ADRA2B*, ρ = 0.83, p = 0.003), and the observation that *HDAC8* expression correlated with object interaction time to the same degree as *HDAC2* expression, but in the opposite direction (∥ρ∥ = 0.71, p = 0.02, Supp. Fig. 1E-F), suggesting that HDAC8, a shorter class I HDAC with many sequence and structural similarities to HDAC2, may provide a compensatory regulatory mechanism like that previously observed between HDAC1 and HDAC2 (Wang et al., 2005, Cho and Cavalli, 2014).

### Significantly changed Gs and Gi, but not Gq, GPCR transcripts were upregulated and positively correlated with object interaction time

Adenylyl cyclase linked Gs GPCRs *ADORA2A, DRD1, GPR6*, and Gi GPCR *DRD2* transcripts were strongly upregulated (log_2_(FC) > 2.4, p < 0.0001, FDR < 0.3) and positively correlated with object interaction time (ρ ≥ 0.8, p < 0.01) (Fig. 2D-E). Pre-synaptic Gi adrenoreceptors *ADRA2B* and *ADRA2C* expression was also positively correlated with object interaction time (ρ ≥ 0.8, p < 0.01, log_2_(FC) ≤ 0.3, p ≥ 0.06).

## Discussion

The findings in this study support a neurotransmitter modulation model that addresses unanswered questions regarding cortical mechanisms of *HDAC2* inhibition (*HDAC2*i)-enhanced cognition. We identified a select group of upregulated CREB-activated genes that modulate cortical synaptic plasticity, cellular differentiation, forebrain development, and neuronal adenosine, catecholamine, and acetylcholine pathways. Long-term *HDAC2*i directly modulated acetylcholine biosynthesis pathways through increased expression via simple CREB regulation, however, dopamine, norepinephrine, and adenosine pathways were modulated through a newly identified *HDAC2*i-induced CREB | Gs signaling positive feedback mechanism. The observed behavioral results of increased novelty preference, latency of enhanced cognition, and longer object exploration times are consistent with these neurotransmitter modulations.

### Activation of a select group of CREB-regulated genes

Our findings are also consistent with literature suggesting that HDAC inhibitors enhance memory processes by activation of genes regulated by the CREB: CBP transcriptional complex (Korzus et al., 2004; Vecsey et al., 2007; Haettig et al., 2011). However, as we have found in previous studies, global stimulation can activate diverse sets of genes in the cortex compared to the hippocampus (Sanders et al., 2019). Indeed, there was no overlap between the set of previously reported DE CREB-regulated hippocampal genes and our DE CREB-regulated cortical genes.

### Positive feedback loop

Both differential expression (Fig. 2D) and correlation evidence (positive behavioral correlation observed with Gs, but not Gq, GPCRs, Supplemental Fig. 1D) implicate adenylyl cyclase linked GPCRs as players in a GPCR → adenylyl cyclase → cAMP → PKA → CREB feedback loop. Cells with transcriptionally accessible adenylyl cyclase-linked receptors (and Gs > Gi) such as those with pre- or post-synaptic adenosine and/or dopamine receptors would enable this positive feedback loop (Fig. 5B), while cells with transcriptionally repressed Gs GPCRs (potentially through cell-dependent DNA methylation patterns (Miller et al., 2008)) would not (Fig. 5A).

### *HDAC2* ASO-treated rats exhibit behavioral and transcriptional changes consistent with reduced norepinephrine, increased dopamine, and increased acetylcholine

The overall findings in this study are consistent with literature linking increased cholinergic and dopaminergic activity with enhanced cognition (Blokland, 1995; Kaasinen and Rinne, 2002; Ballinger et al., 2016; Cameron et al., 2018). However, the surprising result that the injected rats performed no better than controls on the initial day of behavioral testing suggests that a more nuanced interpretation of the data is warranted.

Since norepinephrine release is also associated with cognitive performance (Berridge and Waterhouse, 2003; Sara, 2009; Chalermpalanupap et al., 2013), this study’s finding of decreased cortical expression of the enzyme responsible for norepinephrine synthesis from dopamine (DBH) implies that some aspects of cognition may have been diminished. Norepinephrine has been found to enhance processing of sensory inputs, arousal, and reaction speed. Thus, it is possible that reduced norepinephrine may be at least partially responsible for the observed delay in novelty preference performance (learning latency).

We did not find a linear relationship between the DE genes and individual performance on the behavioral task. However, we did find correlation between 882 DE genes and the time spent interacting with the objects after the initial test day. Novelty preference co-occurred with these longer object interaction times (despite no linear correlation). Since upregulated dopamine availability may increase behavioral rewards, longer object interaction times are likely attributable to its increase. A contributing effect from reduced norepinephrine is also supported by the correlation between object interaction time and both increased inhibitory pre-synaptic α_2_-adrenergic receptor transcripts and decreased post-synaptic α_1_-adrenergic receptor transcripts.

The specific cognitive effects of increased acetylcholine are more difficult to characterize in our study. However, previous studies suggest that increased acetylcholine may be responsible for cognition-enhancing synaptic plasticity and excitability changes (Rasmusson, 2000; Nakajima et al., 1986) like those observed in our study. Additionally, acetylcholine may have played a role in increased object interaction since *CHAT* correlated strongly with object interaction times. This is consistent with literature identifying acetylcholine as a driver of increased attention (Blokland et al., 1995; Ballinger et al., 2016; Hauser et al., 2019).

Taken together, the behavioral findings support the neurotransmitter modulation model implicated by the cortical RNA-seq results. However, although these findings support *HDAC2i’s* ability to promote long-term cortical plasticity and enhanced cognition in rats, it is important to note that the observed cognitive improvements may coincide with potential negative effects such as increased learning latency in novel environments.

### Therapeutic relevance

The findings in the current study support potential roles for *HDAC2* inhibition in treating disorders characterized by cognitive deficits including AD (Graff et al., 2012; Choubey and Jeyakanthan, 2018) and some forms of autism (Kennedy et al. 2016). The evidence for upregulation of dopaminergic signaling found in this study suggests that *HDAC2* inhibition achieved through long-lasting HDAC2 ASOs may also be beneficial for Parkinson’s disease.

It is important to note that the data in this study reveal persistent cortical gene expression alterations produced by long-term *HDAC2* inhibition rather than potentially short-lived therapeutic effects. These persistent pathway modulations suggest new cognitive therapeutic targets such as *CHAT*, *DBH, VMAT2*, and *SV2C*. The SV2 family of proteins has already been successfully targeted in epilepsy treatments to slow neurotransmitter release for patients with focal seizures (Wood et al., 2020), indicating an important role for vesicular packaging modulation in treatment of brain disorders. Our findings suggest that modulation of vesicular packaging may also be useful for promoting cognition-enhancing shifts in neurotransmitter release.

## Materials and Methods

### Subjects

18 adult male Sprague-Dawley rats (4-5 months old; 350 g ± 50 g) were used for this study. All procedures were performed with Vanderbilt University Institutional Animal Care and Use Committee (IACUC)-approved protocols and conducted in full compliance with the Association for Assessment and Accreditation of Laboratory Animal Care (AAALAC).

### Anti-sense oligonucleotides

Hdac2-ASO1 (5’-CToCoAoCTTTTCGAGGTToCoCTA-3’), Hdac2-ASO2 (5’-AToGoCoAGTTTGAAGTCToGoGTC-3’) and non-targeting ASO (NTA) (5’GToToToTCAAATACACCToToCAT-3’) with phosphorothioate and 2’ MOE modified ASO platforms were received from Ionis Pharmaceuticals.

### In vivo ASO administration

Rats were anesthetized with 2% isoflurane and secured in a stereotaxic frame (David Kopf Instruments). *HDAC2* ASOs were administered to male Sprague-Dawley rats through unilateral intracerebroventricular (ICV) injection into the lateral ventricle (Bregma −0.92 mm A/P, −1.4 mm M/L, −3.4 mm D/V). ASOs were diluted to 80 μg/μl in saline and injected 5 mg/kg into the lateral ventricle at a flow rate of 250 nL/minute. Controls were administered the same volume of either non-targeting ASOs (80 μg/μl in saline) or 100% PBS. After the injection, the needle was kept in place for 5 min., followed by suturing of the incision.

### Novelty preference training and testing

On Day 1, rats were introduced to the first object. On Day 2 and following, the object introduced on the previous day (familiar object) was placed in the same location and a new object was introduced (novel object). Rats were recorded for a 10 min test period, followed by a 30-minute familiarization (learning) period while interacting with the two objects (Sanders et al., 2019). Learning was assessed by calculating the difference in the amount of time spent interacting with the novel object (t_o2_) compared to the learned object (t_o1_), expressed as a fraction of the total time spent interacting with the objects:

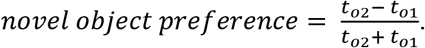

### Tissue collection

Rats were decapitated within 45 minutes of the last behavioral session. Brains were removed and dissected into 15 sections, then flash-frozen for subsequent processing (Sanders et al., 2019).

### Western blots

Tissue from the hippocampus and hippocampal-adjacent cortex were homogenized in RIPA buffer. Protein samples were run on 4-20% TGX Gels (Bio-Rad) and then transferred to PVDF membranes (Millipore) using standard protocols. Primary antibodies were: HDAC2 (Abcam ab12169) and actin (Abcam ab3280). Secondary antibodies were goat anti rat and goat anti rat (Abcam). Membranes were imaged on the LiCor Odyssey fluorescence imaging system.

### RNA extraction from tissue

For rat tissue samples, total RNA was extracted from homogenized left hippocampus and hippocampal-adjacent cortical tissue (Fig. 2A) with AllPrep^®^ DNA/RNA/miRNA kit (Qiagen). Total mRNA was reverse transcribed using the iScript cDNA Synthesis Kit (Bio-Rad). For culture samples the RNeasy plus kit (QIAGEN) and SuperScript VILO (Invitrogen) was used according to manufacturer’s instructions. qPCR was performed with the CFX96 Optical Reaction Module (Bio-Rad) using SYBR green (Bio-Rad). Relative gene expression was determined using the ΔΔCt method (Livak et al., 2010) and normalized to a housekeeping gene.

### Total RNA-seq

Total RNA-seq libraries were prepared from homogenized left hippocampal-adjacent cortical tissue using the TruSeq Stranded Total RNA Library Prep Kit with Ribo-Zero Gold (Illumina) according to manufacturer’s instructions. 1μg of RNA was used as starting material and amplified with 12 PCR cycles. Library size distribution and quality were checked with an Agilent 2100 Bioanalyzer and quantity was determined using qPCR. Samples were verified to have RIN ≥ 8. Libraries were sequenced on Illumina NextSeq instruments using a 75-cycle high throughput kit.

### Statistical analysis, annotation, and visualization

Reads were aligned to the rn5 rat genome and transcriptome using Bowtie2 (Langmead et al., 2012). Differential expression tests were performed using featureCounts (Liao et al., 2014) and edgeR (Robinson et al., 2010) with standard settings. DAVID (Huang et al., 2009) was used for functional annotation of genes.

Normality was formally tested and verified where appropriate. Statistical significance was designated at p < 0.05 for all analyses. Statistical significance was measured using two-sided unpaired t-tests. Adjusted p values were calculated using ANOVA multiple comparisons. FDRs were calculated using Benjamini and Hochberg False Discovery Rate correction.

Bioconductor was used to calculate the most significantly changed transcripts. Changes with ∥log_2_(fold-change)∥ > 0.8, and FDR < 0.3 were considered significant. These thresholds were selected to enable detection of changes close to, or greater than, the observed change in *HDAC2* mRNAs (∥log_2_(fold-change)∥ = 0.9, FDR = 0.28). MATLAB (version R2017; The MathWorks) was used for correlation analysis. The Salk Institute CREB target gene database (Impey et al., 2004, Zhang et al., 2005) was used to identify rattus norvegicus genes with CRE binding domains in the promoter or known enhancers. In a few cases, human data were used to infer likely genes with CREB binding domains. These are indicated in the figures and text.

Gene ontology visualization was performed using Cytoscape (Shannon et al., 2003) with the BINGO plug-in (Maere et al., 2005). Heatmaps were generated with Graphpad Prism version 8 (Graphpad software LLC) and Morpheus (https://software.broadinstitute.org/morpheus).

## Acknowledgments

*HDAC2* ASOs were provided by Ionis pharmaceuticals. Funding was provided by Vanderbilt Pharmacology Department Chair startup funds. Many thanks to Joe Weiss and Ben Coleman for assisting with the rat *HDAC2* ASO intracerebroventricular injections, to Joe Weiss and Casey Paton for assistance with lab work including dissections, Western Blot, and qPCR, to Haley Dotter for quantification of the *HDAC2* ASO-injected rat and control object interaction times, and to Katie Sanders for Bioinformatic graphics support. The author does not have financial or other relationships that constitute a conflict of interest.

## Author Contributions

T.H.S. designed the study and experiments. T.H.S. performed rat injections, behavioral experiments, and RNA-seq analysis. T.H.S. prepared the manuscript.

---

**Supplemental Figure 1.**
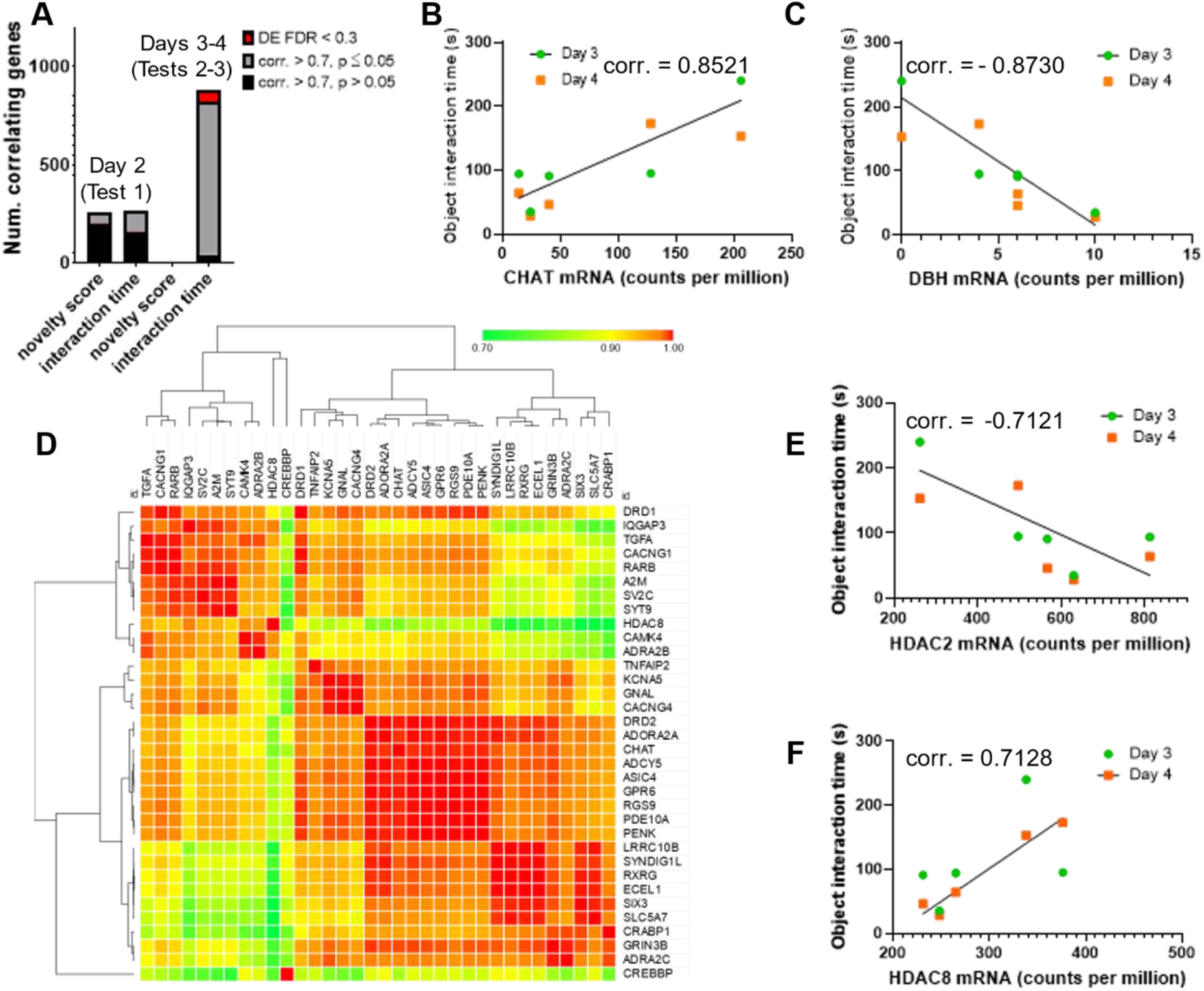
Expression of HDAC and biosynthesis enzyme genes correlated with object interaction time. (**A**) On the first day of testing (Day 2), top differentially expressed (DE, FDR < 0.3) genes did not correlate with behavioral measures (novel object preference or object interaction time). However, 882 genes, including *HDAC2*, correlated with object interaction time on subsequent test days (Days 3 and 4). (**B**) Expression of neurotransmitter biosynthesis enzymes (Fig. 3), *CHAT* and (**C**) *DBH*, correlated significantly with object interaction time. (**D**) Intra-correlation analysis of the top DE genes that correlated with behavior revealed 4 clusters of genes: 1) CREB-regulated genes outside the top group of DE genes, 2) CREB binding protein (*CREBBP*) and *HDAC8*, 3) the top group of CREB-responsive GPCR genes (*ADORA2A, DRD1, DRD2*, and *GPR6*) along with adenylyl cyclase (*ADCY5*) and *CHAT*, and 4) DE genes associated with cell differentiation (*RXRG, SIX3, CRABP1*) and synaptic organization. (**E**) *HDAC2* expression correlated inversely with object interaction time. (**F**) *HDAC8* expression correlated to the same degree as *HDAC2*, but in the opposite direction.

